# Potentially Prebiotic Isocyanide Activation Chemistry Drives RNA Assembly via both Nonenzymatic and Ribozyme-catalyzed Ligation

**DOI:** 10.1101/2025.07.11.664453

**Authors:** Stephanie J. Zhang, Xiwen Jia, Saurja DasGupta, Jack W. Szostak

## Abstract

Nonenzymatic assembly of activated RNA building blocks, such as phosphorimidazolides, would have been essential for the emergence of ribozymes on the early Earth. We previously showed that ribonucleoside monophosphates can be activated to phosphorimidazolides via a prebiotically relevant phospho-Passerini reaction involving 2-aminoimidazole, 2-methylbutyraldehyde, and methyl isocyanide, and that these activated nucleotides enable template-directed nonenzymatic RNA polymerization in the same reaction mixture. Here, we demonstrate that the same chemistry activates oligoribonucleotides and drives both nonenzymatic and ribozyme-catalyzed RNA ligation within the same reaction environment. By demonstrating a continuous path from prebiotic activation chemistry to RNA template copying by both nonenzymatic and ribozyme-catalyzed ligation, our results provide a more integrated and realistic model for RNA assembly on the early Earth.

---

The RNA World hypothesis posits that early life relied on RNA both as the carrier of genetic information and as a catalyst, in the form of RNA enzymes or ribozymes.^1, 2^ Catalytically active RNAs could have emerged via chemical self-assembly pathways such as ligation^3^ of short oligoribonucleotides. In the early stages, RNA replication may have occurred primarily at the level of such short oligonucleotides, while the longer functional RNAs did not need to be replicated.^4^ The eventual appearance of ribozyme ligases and polymerases would have enhanced the efficiency of RNA copying pathways, enabling the direct replication of longer RNA sequences, including ribozymes. Before such ribozymes emerged, however, template-directed RNA copying likely had to rely on energy-rich activated nucleotides such as phosphorimidazolides. In earlier work, we demonstrated that *in situ* nucleotide activation, using a potentially prebiotic phospho-Passerini chemistry based on methyl isocyanide (MeNC), 2-aminoimidazole (2AI), and 2-methyl-butyraldehyde (2MBA), can drive nonenzymatic RNA copying via template-directed polymerization.^5, 6^ This chemistry, likely to occur in UV-irradiated environments rich in ferrocyanide and nitroprusside, provides a direct link between monomer activation and polymerization and could potentially support continuous RNA copying under prebiotic conditions. However, the high concentration of 2AI necessary for nucleotide activation prevents the accumulation of the 5′-5′ imidazolium-bridged dinucleotide intermediate required for nonenzymatic RNA polymerization.^7^ To overcome this problem, we used ice-eutectic conditions, which locally concentrate the reactants and thereby increase the effective concentration of 2AI.^8^ This allowed us to drive nucleotide activation with stoichiometric 2AI while enabling the subsequent formation of the 5′-5′ imidazolium-bridged intermediate. Unlike polymerization, nonenzymatic ligation of 2AI-activated oligonucleotides does not rely on a 5′-5′ bridged intermediate and is therefore not inhibited by excess 2AI.^7, 9^ Although slower than polymerization,^10-12^ ligation better accommodates structured or A/U-rich templates^13^ and proceeds with higher fidelity,^14^ while requiring fewer condensation steps.^15^ *In situ* activation of short oligonucleotides (≤4 nt) has been shown to support polymerization by forming bridged intermediates with monomers.^9^ However, ligation driven by the *in situ* activation of oligonucleotides has not been demonstrated.

The emergence of ribozymes marked a key transition in the origin of life. We previously used *in vitro* selection to identify ligase ribozymes that use 2-aminoimidazole-activated oligoribonucleotides as substrates, thus bridging nonenzymatic and enzymatic RNA ligation.^16^ However, the hydrolytic instability of phosphorimidazolide substrates limits their accumulation, posing a major challenge to uninterrupted RNA assembly. Within protocells, substrate activation would support both chemical and ribozyme-catalyzed RNA ligation, and would have been essential for maintaining effective substrate concentrations and mitigating the inhibitory effects of unactivated substrates.^17^ Here, we outline a prebiotically relevant scenario in which *in situ* activation of RNA substrates enables both nonenzymatic and ribozyme-catalyzed RNA ligation within a shared reaction environment (Fig. 1). We demonstrate that ribozymes remain functional under conditions where RNA substrates are activated, and that substrate activation, nonenzymatic RNA ligation, and ribozyme-catalyzed RNA ligation can all occur in the same environment. Our results support a model for RNA replication on the early Earth, where nonenzymatic RNA assembly – driven by continuous *in situ* activation of its substrates – generated ribozymes that accelerated RNA assembly under the same conditions, paving the way for the rapid propagation of RNA sequences.

**Fig. 1.**
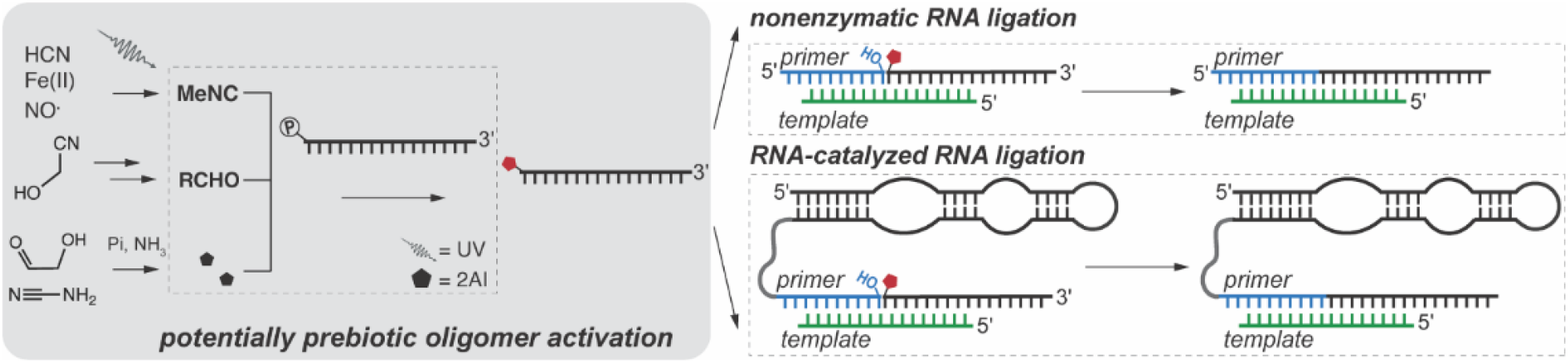
Prebiotic RNA copying *via* templated ligation of activated oligonucleotides generated through potentially prebiotic pathways. The left panel shows a potentially prebiotic pathway for oligoribonucleotide activation using methyl isocyanide, aldehyde, and imidazole. The right panels illustrate two RNA ligation processes: (1) nonenzymatic ligation, where the activated oligonucleotide is chemically ligated to a primer aligned on a template and (2) ribozyme-catalyzed ligation, where a ribozyme facilitates the templated ligation of an activated oligonucleotide to the primer.

Having previously demonstrated the activation of mononucleotides and short oligonucleotides by phospho-Passerini chemistry (henceforth, isocyanide chemistry) in prior work, we sought to first optimize the activation of longer oligonucleotides, which generally show reduced activation efficiency.^5, 6, 9^ Given that longer oligonucleotides would have likely been present at lower concentrations than shorter oligonucleotides and mononucleotides in prebiotic settings,^18, 19^ we began by incubating 10 μM of a 16-nt 5′-monophosphorylated oligoribonucleotide substrate (henceforth, ligator; Table S2, ESI†) with 50 mM MeNC, 50 mM 2MBA, 50 mM 2AI, and 30 mM Mg^2+^ in 200 mM HEPES at pH 8. High-performance liquid chromatography (HPLC) analysis revealed 25.3 ± 0.2% conversion of the 5′-monophosphorylated ligator to its 5′-phosphor-2-aminoimidazolide form after 6 hours (Fig. S1A, ESI†). Extending the reaction to 24 hours did not significantly increase the activation yield, suggesting that the reaction had plateaued by 6 hours (Fig. S1B, ESI†). These conditions served as a baseline for further optimization. Unlike nonenzymatic polymerization, ligation does not require 5′-5′ imidazolium-bridged intermediates, which allowed us to raise 2AI concentrations. After optimization, we achieved >90% activation using 250 mM 2AI and either 250 or 500 mM MeNC and 2MBA (Table 1, reactions 6 and 8). Reducing the concentrations of MeNC or 2MBA below 250 mM caused a marked drop in activation yields, underscoring the need for excess aldehyde and isocyanide, consistent with prior studies on mononucleotide activation.^5, 8, 9^ Interestingly, further increasing 2AI concentrations by 5- to 10-fold led to lower activation yields (Table 1, reactions 3 and 4). In prebiotic environments, 2AI levels could have been limited by UV-induced degradation.^20^ Our results indicate that concentrations of hundreds of millimolar for MeNC and 2MBA, and tens to hundreds of millimolar for 2AI are required for the efficient activation of substrates ranging from monomers to 16-nt oligomers (Tables 1, S1, ESI†). Thus, prebiotic environments capable of sustaining such a range of reagent concentrations could have supported continuous RNA polymerization and ligation by maintaining a diverse pool of activated nucleotides and oligonucleotides.

**Table 1.**
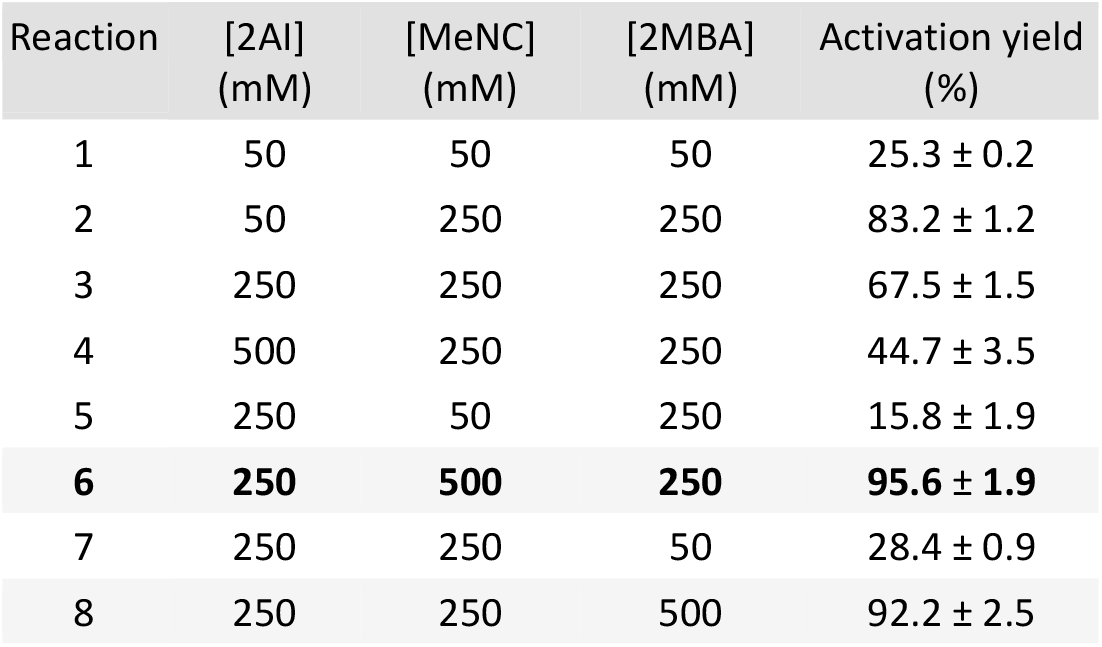
Oligonucleotide activation yields under various conditions as determined by HPLC analysis. Reactions were carried out with 10 μM 5*’*-phosphorylated ligator in 200 mM HEPES at pH 8, 30 mM Mg^2+^, and varying concentrations of 2AI, 2MBA, and MeNC. Samples were incubated at room temperature for 6 hours and then quenched (the optimal time point as identified in Fig. S1, ESI†). Conditions yielding >90% activation are shaded in gray, with the optimal condition (reaction condition 6) highlighted in bold. Error values represent the standard deviation of the mean, based on at least two replicates (n ≥ 2).

We used the optimal condition of 250 mM 2AI, 500 mM MeNC, and 250 mM 2MBA (Table 1, condition 6) to generate activated substrates for nonenzymatic RNA ligation (Fig. 2). Ligators were subjected to activation conditions for 6 hours and subsequently added to the primer-template duplex for ligation without further purification (Fig. 2A). Ligation yields after 3 hours with isocyanide-activated ligators or purified, pre-activated ligators generated *via* EDC coupling were comparable (∼1.5% vs ∼2%) (Fig. S2 and S3), reaching approximately 26% ligation after 24 hours (Fig. 2C). To better simulate prebiotic conditions, where no artificial boundaries would exist between substrate activation and RNA copying, we tested whether oligonucleotide activation and nonenzymatic ligation could proceed in the same pot (Fig. 2B). Upon adding an unactivated ligator to the primer-template complex under activation chemistry, only ∼7% of the primer was ligated after 24 hours (Figs. 2C, S3). A previous study has shown that 2AI-activated ligators undergo minimal hydrolysis under similar conditions over similar timescales,^16^ suggesting that hydrolysis was not responsible for the reduced yield. Based on previous findings, the reduced ligation yields likely stem from poor activation of the ligator due to steric hindrance at the ligation junction duplex, which makes its 5′-phosphate group less accessible to activating reagents. If so, a fluctuating environment that allows for oligonucleotides to repeatedly bind to and dissociate from template strands could lead to increased substrate activation and ligation (see later).

**Fig. 2.**
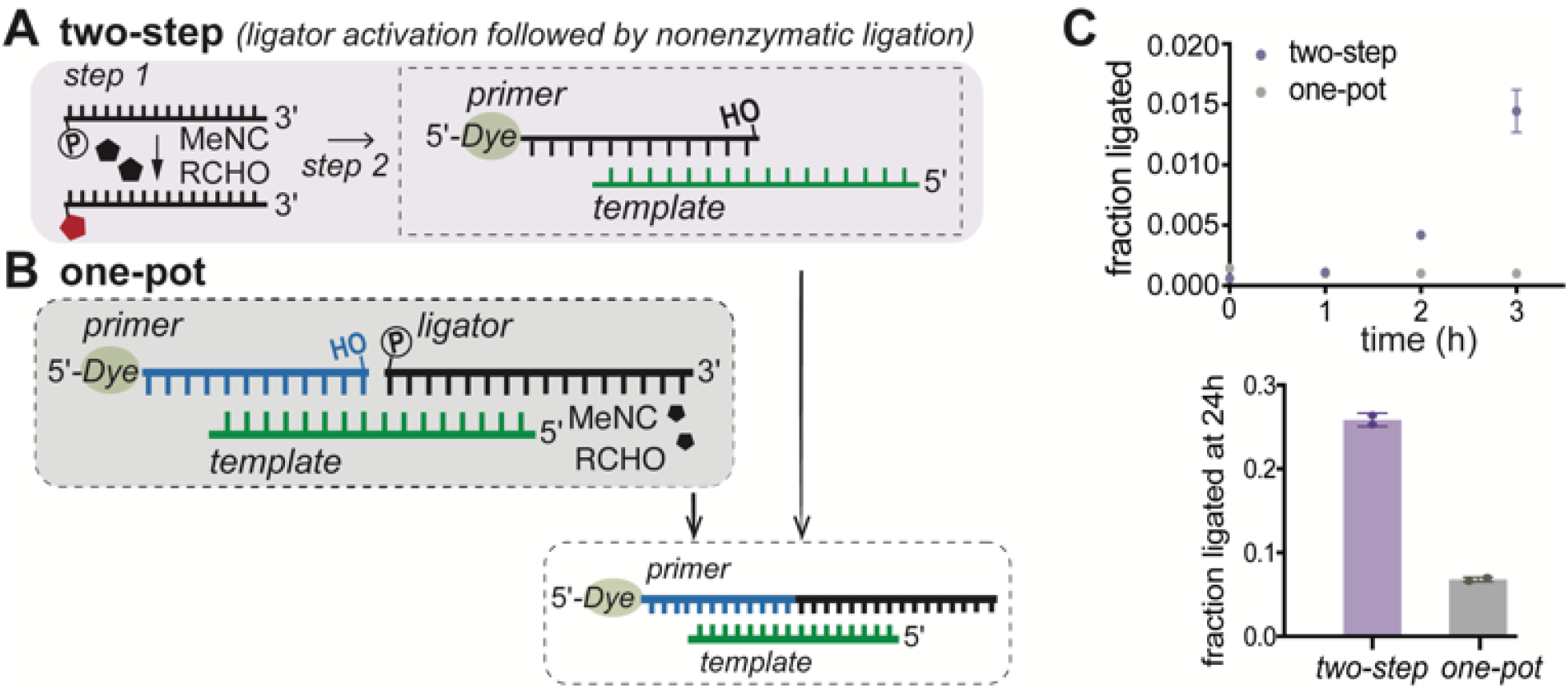
Nonenzymatic RNA ligation driven by prebiotic activation chemistry. Schematic of nonenzymatic ligation using **(A)** ligators activated *via* isocyanide activation chemistry prior to the ligation reaction, and **(B)** ligators activated *in situ* during the ligation. **(C)** Time-course analysis of nonenzymatic ligation with ligators either pre-activated (purple) or activated *in situ* (gray) with isocyanide chemistry. The right inset compares ligation yields after 24 hours. Ligation reactions were analyzed by denaturing PAGE (see Fig. S3, ESI†), containing 1.2 μM primer, 1.32 μM template, 200 mM HEPES (pH 8), 50 mM Mg^2+^, and 2.4 μM of either the isocyanide-activated ligator mixture (added after 6 hours of activation) or activated directly within the ligation reaction. The increase in the rate of ligation over time likely results from the gradual increase in the concentration of activated ligators.

We next examined whether the same activation chemistry could also support ribozyme-catalyzed RNA ligation. We began by assessing the compatibility of ligase ribozyme activity with the conditions of chemical activation. To do this, we introduced 500 mM MeNC, 250 mM 2MBA, and 250 mM 2AI into a ribozyme-catalyzed ligation reaction using pre-activated, purified ligators in 200 mM HEPES at pH 8 and 10 mM Mg^2+^. We observed a decrease in ligase activity in the presence of 2AI (Fig. S4, ESI†), with a marked reduction in ligation yield above 50 mM 2AI (Fig. S5A, ESI†). To test if 2AI interferes with ligation by chelating Mg^2+^ and consequently reducing its availability for RNA catalysis, we used a fluorescence assay^21^ based on the Mg^2+^-sensitive dye (Mag-fura-2) to quantify free Mg^2+^. The assay revealed a progressive decrease in free Mg^2+^ concentration as 2AI levels increased, with detectable effects at 2AI concentrations as low as 5 mM (Fig. S6, ESI†). To counteract this effect, we increased the Mg^2+^ concentration from 10 mM to 200 mM in reactions containing 200 mM 2AI. This adjustment restored ribozyme activity, increasing ligation yield from 4.1 ± 0.1% to 33.3 ± 0.1% after 3 hours (Fig. S5B, ESI†). As this ribozyme reaches catalytic saturation at 4 mM Mg^2+^,^22^ the observed improvement is likely due to the added Mg^2+^ restoring the free Mg^2+^ pool in the presence of 2AI.

Having confirmed that ligase ribozyme function is compatible with the optimized activation conditions, we integrated the isocyanide activation chemistry with ribozyme-catalyzed ligation (Figs. 3, S7, ESI†). Unactivated ligators were activated for 6 hours and then added to the ribozyme primer-template duplex without any purification, yielding 42 ± 1% ligation after 3 hours, which is comparable to yields obtained with purified ligators pre-activated with EDC (Fig. S5A, ESI†). Extending the reaction time to 24 hours increased the yield to 54 ± 1% (Figs. 3B, S7A, ESI†), demonstrating that oligonucleotides activated by potentially prebiotic chemistry can drive efficient ribozyme-catalyzed ligation. We then tested whether substrate activation and ribozyme-catalyzed ligation can proceed in a shared environment, mimicking a prebiotic scenario with no separation between chemical activation and RNA copying by ribozymes. We added the ribozyme primer-template duplex, the unactivated ligator, and activation reagents in one pot (Fig. 3C). We observed a gradual activation of the ligator over the first 6 hours (Figs. 3D, S7B, ESI†), and a ligation yield of 23 ± 1% after 24 hours. This was roughly half of that achieved when the ligators were pre-activated by isocyanide chemistry (Figs. 3B, 3D). To test if this decrease was due to inhibition by template-bound unactivated ligators, we incubated the ribozyme-primer construct and template with varying ratios of purified activated and unactivated ligators. We observed that increasing the fraction of unactivated ligators resulted in a marked decrease in ligation, suggesting that unactivated ligators competitively inhibit ligation (Fig. S8, ESI†). This reduced efficiency of the one-pot ribozyme-catalyzed ligation reaction is comparable to that observed for nonenzymatic ligation (Figs. 2C, 3C).

**Fig. 3.**
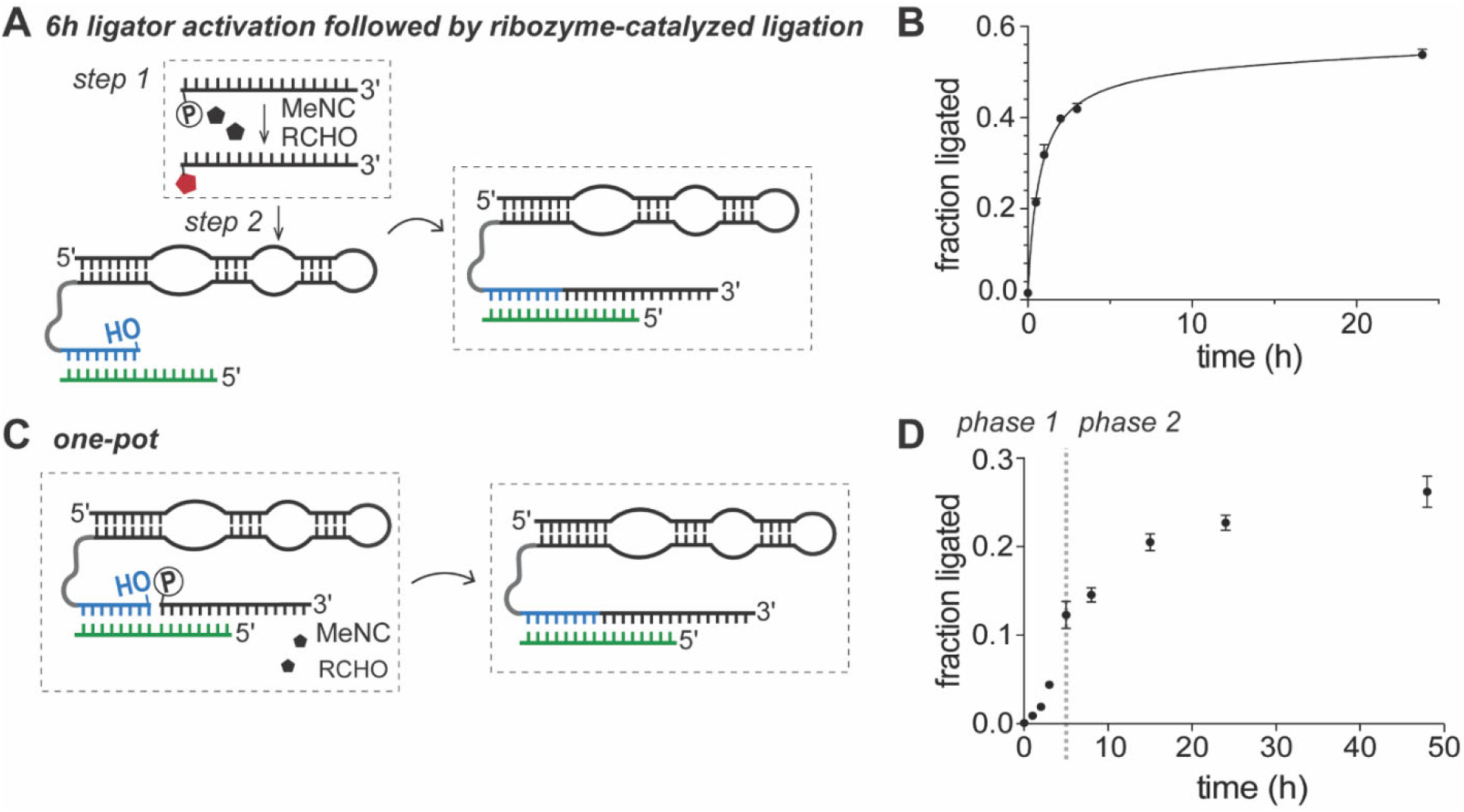
MeNC activation of oligonucleotides drives ribozyme-catalyzed RNA ligation. **(A)** Ribozyme-catalyzed ligation of pre-activated oligonucleotides. The unactivated ligator was incubated under activation conditions for 6 h and then added to a mixture containing the ribozyme-primer construct and the template without any purification. Ligation reactions contained 1.2 μM ribozyme-primer construct, 1.32 μM template, and 2.4 μM isocyanide-activated ligators in the presence of 200 mM HEPES at pH 8 and 50 mM Mg^2+^. **(B)** Time course of ribozyme-catalyzed ligation in (A) (See Fig. S7A, ESI† for the corresponding denaturing PAGE data). **(C)** *In situ* MeNC activation of oligonucleotides enables ribozyme-catalyzed ligation in one pot. 1.2 μM ribozyme-primer construct and 1.32 μM template were incubated with 2.4 μM unactivated ligators in the presence of 250 mM MeNC, 250 mM 2MBA, 250 mM 2AI, 200 mM HEPES at pH 8, and 50 mM Mg^2+^. **(D)** Time course of ribozyme-catalyzed ligation in (C) (See Fig. S7B, ESI† for the corresponding denaturing PAGE data). The plot is biphasic: phase 1 reflects increasing ligation as activated ligators accumulate. In phase 2, ligation proceeds more rapidly but slows as the reaction approaches a plateau.

We hypothesized that the reduced activation efficiency in one-pot reactions was due to the limited accessibility of the activation reagents to the 5′-phosphate group of the unactivated ligator, likely sequestered at the ligation junction when bound to the template. To test this possibility, we incubated the ribozyme-primer construct with the unactivated ligator under activation conditions but withheld the template for the first 3 hours (Fig. S9, ESI†). We reasoned that in the absence of template binding and the formation of a tight ligation junction, the ligator 5′-phosphate group should remain more accessible, allowing more efficient activation. As expected, this approach improved ligation yield to 33.2 ± 0.5 % in 24 h. The inhibitory effect of template-bound unactivated ligators is particularly pronounced with longer oligonucleotides, which form stable duplexes that hinder exchange with activated ligators. In a prebiotic scenario, where shorter ligators would be more abundant, the rapid dissociation of unactivated ligators from their templates is expected to favor ribozyme-catalyzed ligation of ligators that have been activated off-template. However, weak ligator-template interactions may have a detrimental effect on ribozyme-catalyzed RNA ligation. To test this possibility, we repeated the ligation reaction with templates shortened by 1-3 nucleotides from their 5′ end (Fig. S10, ESI†). Reducing base-pairing between the template and the ligator from 7 to 5 base pairs had only a modest effect on the ligation yield. Therefore, we suggest that environments with fluctuating temperatures, salt or pH, which allow longer oligos with weak interactions with their templates to dissociate, would favor substrate activation and subsequent ligation by the ribozyme when they are again bound to the template.

Our findings indicate that a high concentration of reagents, especially, isocyanide is crucial for efficient oligonucleotide activation. The continuous (or periodic) photochemical generation of methyl isocyanide upon UV irradiation (Fig. 1) could enhance ligation by re-activating hydrolyzed unactivated ligators.^4^ Such a mechanism could sustain cycles of RNA activation and copying. We performed ribozyme-catalyzed ligation, where 5′-monophosphorylated (i.e., unactivated) ligators, MeNC, 2MBA, and 2AI were added daily over four consecutive days. This setup mimics a prebiotic scenario in which both oligonucleotides and activating agents are intermittently replenished due to geochemical or photochemical fluctuations. Repeated additions increased ligation yield from 18% on day 1 to 43.7% by day 4 (Fig. 4). Successive additions of activation reagents on the early Earth due to geochemical fluctuations could have enabled comparable oligonucleotide activation, supporting multiple cycles of ribozyme-catalyzed RNA copying.

**Fig. 4.**
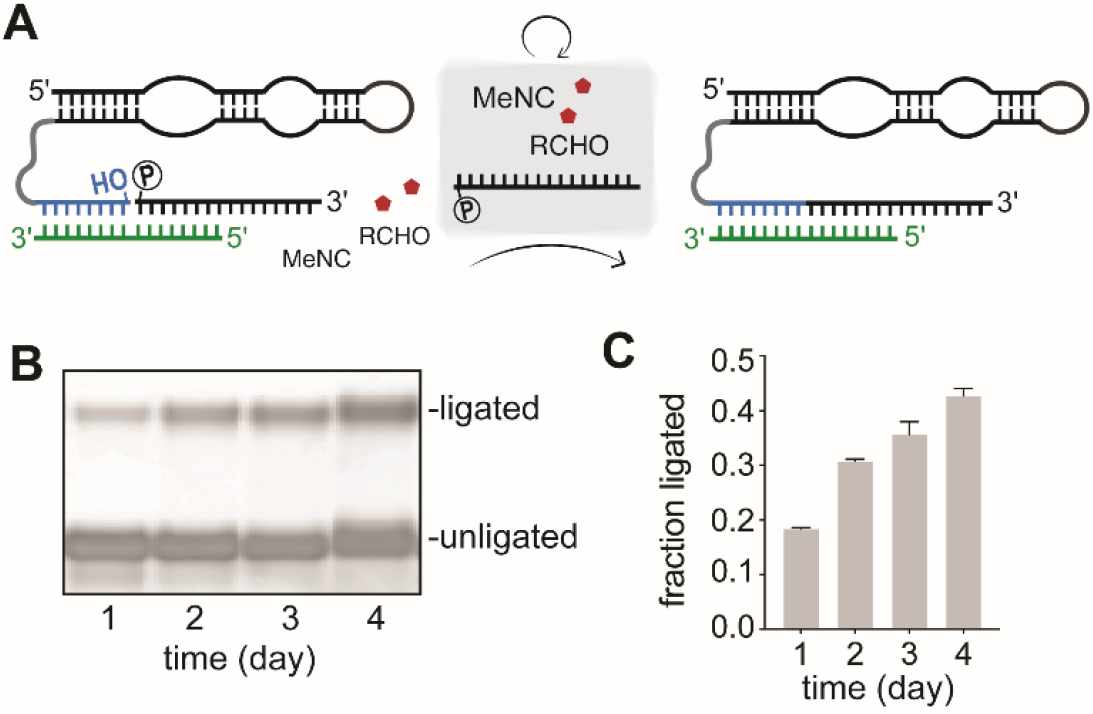
Cycles of activation chemistry drive efficient ribozyme-catalyzed RNA ligation. **(A)** Daily addition of unactivated ligator and activation reagents to the ribozyme-catalyzed ligation reaction increases ligation yield. **(B)** Denaturing PAGE showing increased ribozyme-catalyzed RNA ligation driven by repeated cycles of ligator activation over 4 days. **(C)** Ligation yield after each daily activation cycle over 4 days. Error bars represent standard deviations of the mean, n = 2 replicates.

Recent research has identified a one-pot reaction where 3′-monophosphorylated oligonucleotides are activated by diamidophosphate and imidazole to produce oligonucleotide 2′, 3′-cyclic phosphates. This activation was shown to facilitate hairpin ribozyme-catalyzed RNA ligation.^23^ However, hairpin ribozyme-catalyzed ligations were performed under ice-eutectic conditions to shift the equilibrium in favor of ligation over cleavage. The cyclic phosphate pathway produces ∼30% ligation after 30 days, whereas ligation with 5’-phosphorimidazolides reaches similar yields within 24 – 48 hours. Moreover, ligation with 2′, 3′-cyclic phosphate ligators is extremely slow without enzymes, whereas 5*’*-phosphorimidazolides support both nonenzymatic and ribozyme-catalyzed ligation in the same environment. By demonstrating the activation of oligonucleotide substrates and their nonenzymatic and enzymatic ligation under a unified set of conditions, this work offers a comprehensive experimental model of prebiotic RNA assembly.

While this framework offers insight into how prebiotic activation chemistry could support RNA ligation, several limitations remain. First, although MeNC can be generated under simulated prebiotic conditions, their geochemical availability, stability, and accumulation on the early Earth are not well established. Second, the activation conditions used here involve relatively high concentrations of MeNC, 2AI, and aldehydes.^5, 6^ While such concentrations are useful for modeling these reaction pathways, their prebiotic relevance is uncertain. However, our observation that repeated cycles of reagent and oligonucleotide addition sustain ligation over multiple days suggests that dynamic environments, such as wet-dry cycles, could allow continuous substrate turnover and mitigate inhibitory by-product build-up. Finally, we find that template-bound oligonucleotides are less efficiently activated. This observation implies that environmental fluctuations that promote strand dissociation and reannealing may be critical in prebiotic settings. Such cycles could allow oligonucleotides to be activated in their unbound state and subsequently undergo template-directed ligation under conditions favorable to hybridization.

We are currently attempting to evolve ribozymes that facilitate monomer and oligomer activation, which would make substrate activation and reactivation much more efficient. Ongoing efforts to integrate prebiotic activation chemistries into model protocells, such as lipid vesicles or coacervates,^24, 25^ may reveal how compartmentalization could have influenced early biochemical reactions. In that context, it is important to determine how activation chemistry affects the chemical integrity of protocells. Ultimately, incorporating these mechanisms into flow systems that mimic environmental cycles could help to develop self-replicating protocells capable of supporting potentially prebiotic RNA assembly.

## Author contributions

S.J.Z., X.J., and S.D. designed the research; S.J.Z. and X.J. performed the research; S.J.Z., X.J., S.D., and J.W.S. analyzed the data; and S.J.Z., S.D., and J.W.S. wrote the paper.

J.W.S. is an Investigator of the Howard Hughes Medical Institute. This work was supported in part by grants from the Simons Foundation (290363) and the NSF (2325198) to J.W.S., and by University of Notre Dame Startup Funds to S.D. S.J.Z. would also like to acknowledge support by a postdoctoral fellowship (F32 AG087642) from the National Institutes of Health.

## Data availability

The data supporting this article have been included as part of the ESI.†

