## Supplementary Information for "Potentially Prebiotic Isocyanide Activation Chemistry Drives RNA Assembly via both Nonenzymatic and Ribozyme-catalyzed Ligation"

Electronic Supplementary Information

### CONTENTS


**1. Materials and Methods** **S2–S4**

1.1 General information S2

1.2 Potentially prebiotic activation of oligonucleotides using 2-aminoimidazole S2

1.3 High-performance liquid chromatography (HPLC) analysis of ligator activation S2

1.4 Nonenzymatic RNA ligation reaction and analysis of the ligation products S2

1.5 Ribozyme-catalyzed RNA ligation reaction and analysis of the ligation products S3

1.6 Effect of template length on ribozyme-catalyzed RNA ligation S3

1.7 Measurement of free $Mg^{2+}$ concentration in the presence of 2-aminoimidazole S4

1.8 Ribozyme-catalyzed RNA ligation reactions with cycles of substrate reactivation S4

**2. Supplementary Figures** **S5–S14**

**3. Supplementary Tables** **S15**

**4. References** **S16**

### 1. Materials and Methods

#### 1.1 General information

*Materials.* Reagents and solvents were obtained at the highest purity available from Acros Organics, Alfa Aesar, Fisher Scientific, Sigma-Aldrich, ThermoFisher Scientific, or Tokyo Chemical Industry Co., and were used without any further purification unless otherwise noted. RNA oligonucleotides were purchased from Integrated DNA Technologies. All reactions were carried out in UltraPure$^{TM}$ DNase/RNase-free distilled water.

*Preparation and storage of stock solutions*. Stock solutions of 2-aminoimidazole hydrochloride and buffers such as 2-[4-(2-hydroxyethyl)piperazin-1-yl]ethanesulfonic acid (HEPES) were prepared by dissolving the corresponding reagent in UltraPure$^{TM}$ DNase/RNase-free distilled water. After adjusting the pH to the reported values with NaOH or HCl, the stock solution was filter-sterilized with 0.22 µm syringe filters (Millipore Sigma). Each stock solution was then aliquoted and kept at -20 °C until further use.

#### 1.2 Potentially prebiotic activation of oligonucleotides using 2-aminoimidazole

Each activation reaction (100 µL) contained 10 µM ligator, the indicated concentrations of 2-aminoimidazole (2AI), methyl isocyanide (MeNC), 2-methylbutyraldehyde (2MBA), 30 mM $MgCl_2$, and 200 mM HEPES at pH 8.0. The solution was briefly vortexed, ensuring proper mixing. The reaction was incubated at room temperature for 6 h, which was the optimal incubation time as determined by the time course (Fig. S1B). The progress of the reaction was monitored by reverse phase high-performance liquid chromatography (HPLC) over the course of 24 h (see details in section 1.3). The kinetics of the reaction was followed by quantifying peak areas corresponding to the unactivated and activated ligators at different time points.

#### 1.3 High-performance liquid chromatography (HPLC) analysis of ligator activation

Each activation reaction (100 µL) containing 10 µM ligator, the indicated concentrations of 2AI, MeNC, 2MBA, 30 mM $MgCl_2$, and 200 mM HEPES at pH 8.0 was incubated for 6 h. The activation reagents were removed using a ZYMO Oligo Clean & Concentrator spin column (ZYMO Research). The unreacted 5'-phosphorylated ligator and 2AI-activated ligator were separated and analyzed on an Agilent 1100 series HPLC system equipped with a solvent degasser, an autosampler, and a fraction collector. The sample was separated on a 4.6 mm x 260 mm, 5 µm Eclipse Plus C18 column (Agilent Technologies) using gradient elution between (A) aqueous 25 mM triethylamine/bicarbonate buffer at pH 7.5 and (B) acetonitrile, where the sample was eluted between 5 % and 11 % B over 40 min. Fractions containing unactivated and activated ligators were flash-frozen and lyophilized before confirming their identities by liquid chromatography-mass spectrometry (LC-MS). Specifically, the lyophilized samples were resuspended in Milli-Q water and analyzed on an Agilent 1200 HPLC coupled to an Agilent 6230 TOF equipped with a solvent degasser, column oven, autosampler, and diode array detector. The sample was separated by IP-RP-HPLC on a 100 mm × 1 mm (length × i.d.) Xbridge C18 column with 3.5 µm particle size (Waters Corporation) using gradient elution between (A) aqueous 200 mM 1,1,1,3,3,3-hexafluoro-2-propanol with 1.25 mM triethylamine, pH 7.0, and (B) methanol, where the sample was eluted between 2.5 % and 20 % B over 28.5 min with a flow rate of 125 µL/min at 60 °C. Samples were analyzed in negative mode from 239 m/z to 3200 m/z with a scan rate of 1 spectrum/s and the following instrument settings: drying gas flow, 8 L/min; drying gas temperature, 325 °C; nebulizer pressure, 30 psig; capillary voltage, 3500 V; fragmentor, 200 V; and skimmer, 65 V.

#### 1.4 Nonenzymatic RNA ligation reaction and analysis of the ligation products

Typical experiments use a fluorescent dye-labeled primer, and ligation products are characterized by denaturing polyacrylamide gel electrophoresis (PAGE), where different primer extension lengths yield distinct banding patterns. However, many dyes have functional groups, such as carboxyls and amines, which are incompatible with isocyanide activation chemistry. As described previously,[1] we employed a post-labeling strategy, introducing the dye after the reactions were complete. We used a thiol-modified primer, validated to be unreactive to isocyanide activation chemistry, enabling subsequent conjugation of a maleimide-fluorophore via sulfhydryl-maleimide coupling.

2.67 μM thiol-modified primer and 2.94 μM template RNAs were annealed in a 67.3 μL solution containing 445.6 mM HEPES at pH 8.0 by heating at 90 °C for 2 min and then cooled to 23 °C at a rate of 0.1 °C/s. The reaction was initiated by the addition of 82.7 μL containing either activated ligator or the unactivated ligator with activation reagents (2AI, 2MBA, and MeNC) and $MgCl_2$, resulting in a 150 μL reaction volume with final concentrations of 1.2 μM primer, 1.32 μM template, 2.4 μM ligator, 250 mM 2AI, 250 mM 2MBA, 500 mM MeNC, 200 mM HEPES, and 50 mM $MgCl_2$.

Upon reaction completion, the reaction mixture was resuspended in 30 μL 100 mM HEPES at pH 7.5, and disulfide bonds were reduced using a 10-fold molar excess of tris-(2-carboxyethyl) phosphine hydrochloride (TCEP). 1 mM Alexa 488 C5 maleimide in anhydrous dimethyl sulfoxide (DMSO) was added to the primer-template duplex dropwise, and the reaction was allowed to proceed at room temperature for 2 h, protected from light. The labeled primer-template duplex was separated from free dye using Oligo Clean & Concentrator spin columns (ZYMO Research), resuspended in 5 μL of 100 mM HEPES buffer, and mixed with 30 μL of quenching buffer containing 8.3 M urea, 1.3× Tris/Borate/EDTA (TBE) buffer at pH 8.0, 0.005 % Bromophenol Blue, and 0.04 % Orange G. The primers, templates, and ligators used in the ligation assays are listed in Table S2.

Denaturing polyacrylamide gels were prepared using the SequaGel–UreaGel system (National Diagnostics). The gels were allowed to pre-run at a constant power of 20 W for 40−45 min. Aliquots (15 μL each) of samples were separated by 20 % (19:1) denaturing PAGE at a constant power of 5 W for 10 min (to facilitate uniform sample entry and initial band alignment) and then at 25 W for 1 h. The gels were imaged on a Typhoon 9410 scanner (GE Healthcare) and quantified using the accompanying ImageQuant TL software. All reactions were performed in duplicate or greater.

#### 1.5 Ribozyme-catalyzed RNA ligation reaction and analysis of the ligation products

2.67 μM ribozyme-primer and 2.94 μM template RNAs were annealed in a 67.3 μL solution containing 445.6 mM HEPES at pH 8.0 by heating at 90 °C for 2 min and then cooled to 23 °C at a rate of 0.1 °C/s. The reaction was initiated by the addition of 82.7 μL containing either activated ligator or the unactivated ligator with activation reagents (2AI, 2MBA, and MeNC) and $MgCl_2$, resulting in a 150 μL reaction volume with final concentrations of 1.2 μM ribozyme-primer, 1.32 μM template, 2.4 μM ligator, 250 mM 2AI, 250 mM 2MBA, 500 mM MeNC, 200 mM HEPES, and 50 mM $MgCl_2$.

Aliquots (15 μL each) were removed at given time points and desalted using Oligo Clean & Concentrator spin columns (ZYMO Research). The isolated material was resuspended in 5 μL of 100 mM HEPES buffer and mixed with 30 μL of quenching buffer containing 8.3 M urea, 1.3× TBE buffer at pH 8.0, 0.005 % Bromophenol Blue, and 0.04 % Orange G. The ligators, ribozyme-primer constructs and templates used in the ligation assays are listed in Table S2.

10 % denaturing polyacrylamide gels were prepared using the SequaGel–UreaGel system (National Diagnostics). The gels were allowed to pre-run at a constant power of 20 W for 40−45 min. 15 μL aliquots of samples were separated by 10 % (19:1) denaturing PAGE at a constant power of 5 W for 10 min and 25 W for 1 h. The gels were incubated with 1X SYBR® Gold stain for 5 min, imaged on a Typhoon 9410 scanner (GE Healthcare), and quantified using the accompanying ImageQuant TL software. All reactions were performed in duplicate or greater.

#### 1.6 Effect of template length on ribozyme-catalyzed RNA ligation

The ribozyme-primer and template were first annealed in a 12.1 μL solution containing 3 μM ribozyme-primer and 3.3 μM template (comprising four different lengths: template (16 nt), template-1 (15 nt), template-2 (14 nt), and template-3 (13 nt), see Table S2), along with 494.6 mM HEPES at pH 8. The annealing process involved heating at 90 °C for 2 min, followed by cooling to 23 °C at a rate of 0.1 °C/s. Subsequently, 1.5 µL of $MgCl_2$ was added to the stock solution, and the construct was refolded at 55 °C for 10 min, followed by cooling to 23 °C at a rate of 0.1 °C/s. This was followed by the addition of 4.4 µM unactivated ligator, 458 mM HEPES at pH 8.0, 366.5 mM 2AI at pH 8.0, 458.2 mM 2MBA, and 916.3 mM MeNC to initiate the reaction. Finally, the solutions were quenched after 24 hours using Oligo Clean & Concentrator spin columns (ZYMO Research).

#### 1.7 Measurement of free $Mg^{2+}$ concentration in the presence of 2-aminoimidazole

Free $Mg^{2+}$ concentration was measured using the $Mg^{2+}$-sensitive dye Mag-fura-2 (Invitrogen) following a previously reported procedure.[2] Fluorescence measurements were conducted on a Cary Eclipse Spectrophotometer (Varian), with the emission wavelength fixed at 500 nm and excitation wavelengths ranging from 300 to 400 nm. A standard curve correlating the fluorescence intensity ratio ($I_{340\ nm}/I_{370\ nm}$) to free $Mg^{2+}$ concentration was generated (Fig. S6A). Standard curve conditions included 5 µM Mag-fura-2, 200 mM HEPES at pH 8.0, and $MgCl_2$ (0-14 mM). The fluorescence properties of Mag-fura-2 were minimally affected by 2AI (Fig. S6B). To assess the effect of 2AI on free $Mg^{2+}$ levels, we varied either the concentrations of 2AI (0-14 mM) in the presence of constant $Mg^{2+}$ (14 mM) (Fig. S6C), or the concentrations of $Mg^{2+}$ (0-14 mM) in the presence of constant 2AI (14 mM) (Fig. S6D). The standard curve (Fig. S6A) was used to estimate free $Mg^{2+}$ levels in these conditions (Figs. S6C–D). Reaction mixtures for testing the effect of 2AI on Mag-fura-2 included 5 µM Mag-fura-2, 200 mM HEPES at pH 8.0, and the indicated concentration of 2AI at pH 8.0.

To estimate the dissociation constant ($K_D$) between 2AI and $Mg^{2+}$, we quantified the concentration of free $Mg^{2+}$ in solutions containing a fixed total $Mg^{2+}$ concentration (14 mM) and increasing concentrations of 2AI, as measured by the Mag-fura-2 fluorescent assay (Fig. S6C). Each data point represents the mean of triplicate measurements, with standard deviations used to weight the fit. We modeled $Mg^{2+}$–2AI binding using a simple 1:1 reversible equilibrium binding model:

$$[Mg^{2+}]_{free} = \frac{K_D \cdot [Mg^{2+}]_{total}}{Kd + [2AI][Mg^{2+}]_{free}}$$

This model assumes reversible complexation between one molecule of 2AI and one $Mg^{2+}$ ion, where the observed free $Mg^{2+}$ is determined by the total $Mg^{2+}$ and the concentration of free 2AI. The curve was fitted using non-linear least squares regression (Python, scipy.optimize.curve_fit), with the full analysis script available in our OSF repository.[3] This fit yielded an apparent $K_D$ of 41.5 $\pm$ 0.6 mM. However, this value exceeds the dynamic range of the Mag-fura-2 assay, which approaches saturation near 14 mM $Mg^{2+}$. As such, the extrapolated $K_D$ should be interpreted with caution. The minimal change in fluorescence signal across the titration supports the conclusion that 2AI binds $Mg^{2+}$ only weakly, but the precise $K_D$ remains uncertain due to assay limitations.

#### 1.8 Ribozyme-catalyzed RNA ligation reactions with daily cycles of substrate activation

The *in situ* activation-driven ribozyme-catalyzed ligation reaction was set up as outlined in section 1.5. A fresh batch of unactivated ligators (to 2.4 µM) was added to the ligation reaction in the presence of 250 mM 2MBA, 200 mM 2AI, and 500 mM MeNC, once daily for 4 days. The ligation reaction mixture was then quenched and desalted at specified time points (days 1–4) using Oligo Clean & Concentrator spin columns (ZYMO Research).

### 2. Supplementary Figures

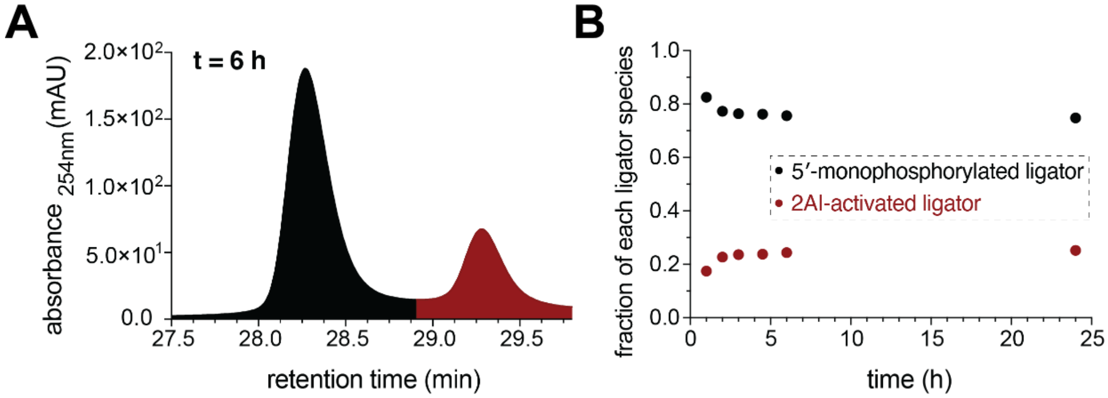


**Fig. S1 Potentially prebiotic activation of a 16-nt RNA ligator with 2AI in the presence of 2MBA and MeNC. (A)** HPLC trace at t = 6 h showing peaks at 28.4 and 29.4 min corresponding to 5′-monophosphorylated ligator and 2AI-activated ligator, respectively. **(B)** Time course for ligator activation with 2AI. Quantification of the 5′-monophosphorylated ligator and 2AI-activated ligator fractions by HPLC as a function of time. Reaction condition: 10 µM ligator, 50 mM 2AI, 50 mM MeNC, 50 mM 2MBA, 200 mM HEPES at pH 8, and 30 mM $MgCl_2$ at room temperature.

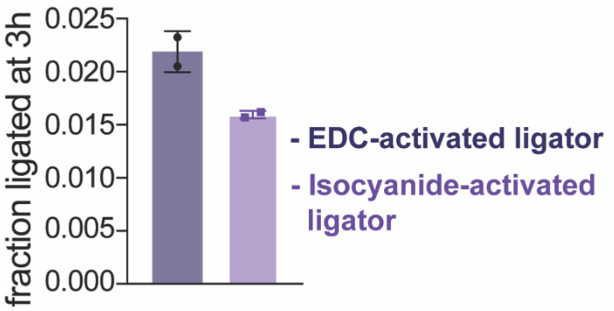


**Fig. S2 Ligation yields with isocyanide-activated and unpurified or EDC-activated and purified ligators.** Ligation yields after 3 hours using either pre-activated ligators prepared using EDC coupling and purified by HPLC as control, or unpurified ligators activated *via* isocyanide chemistry. Ligation reactions were carried out with 1.2 µM primer, 1.32 µM template, and 2.4 µM of either purified, pre-activated ligators using EDC coupling, or ligators activated by isocyanide chemistry (added after 6 hours of activation without purification), in the presence of 200 mM HEPES at pH 8 and 50 mM $MgCl_2$.

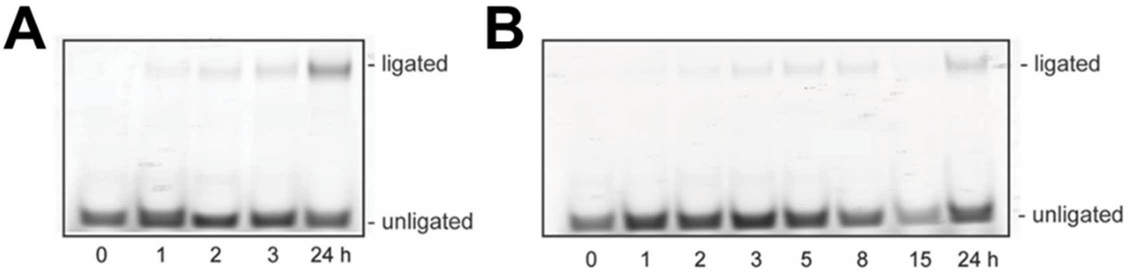


**Fig. S3 Denaturing PAGE depicting the time-course of nonenzymatic RNA ligation using pre-activated or *in situ* activated ligators.** Denaturing PAGE corresponding to the data presented in Fig. 2D. Products of nonenzymatic ligation using **(A)** unpurified isocyanide-activated ligators (added after 6 h of isocyanide activation) or **(B)** ligators activated *in situ* were assayed by denaturing PAGE. Reaction conditions: (A) Primer and template were added to 2.4 µM ligators pre-activated by isocyanide chemistry (250 mM MeNC, 500 mM 2MBA, 250 mM 2AI) to final concentrations of 1.2 µM and 1.32 µM, respectively, in the presence of 200 mM HEPES at pH 8 and 50 mM $MgCl_2$, (B) 1.2 µM primer, 1.32 µM template and 2.4 µM unactivated ligator were incubated with 200 mM HEPES at pH 8 and 50 mM $MgCl_2$ in the presence of 250 mM MeNC, 500 mM 2MBA, and 250 mM 2AI.

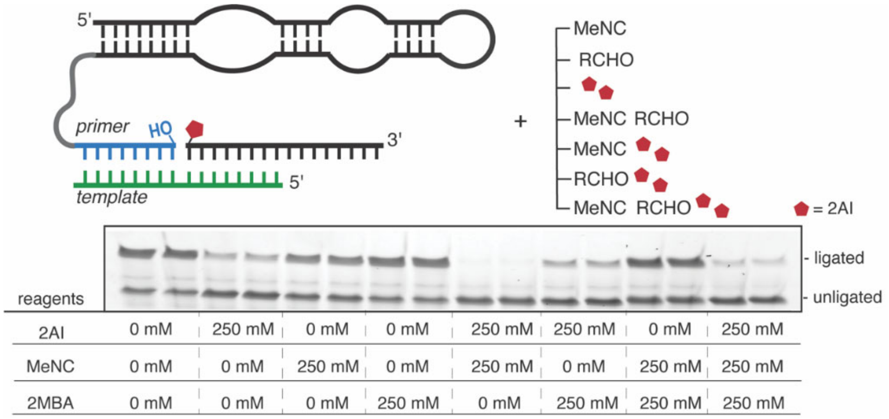


**Fig. S4 Inhibition of ribozyme-catalyzed RNA ligation under conditions of activation chemistry is primarily due to 2AI.** The indicated amounts of activation reagents were added to a ribozyme-catalyzed ligation reaction containing 1.2 µM ribozyme-primer construct, 1.32 µM template, 2.4 µM pre-activated ligator, 200 mM HEPES at pH 8, and 10 mM $MgCl_2$. The addition of MeNC or 2MBA or both had an insignificant effect on ligation; however, the presence of 2AI caused a marked decrease in ligation, either alone or in combination with other reagents. Each condition was assayed in duplicate.

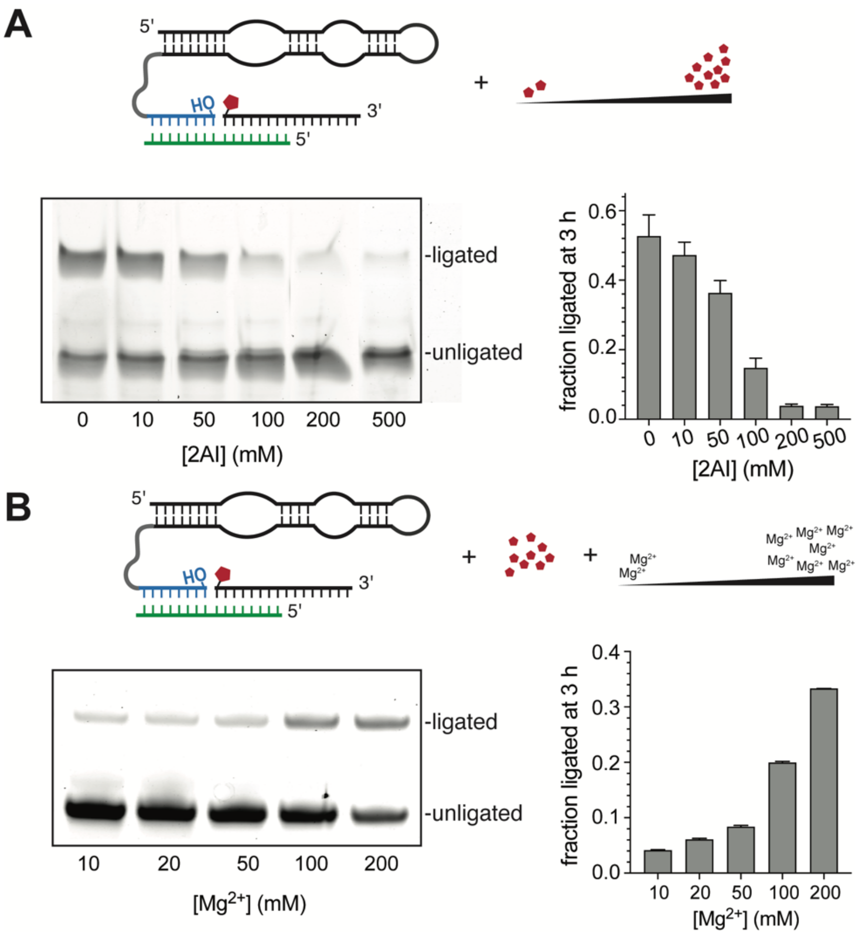


**Fig. S5 Increasing $Mg^{2+}$ concentration rescues ribozyme-catalyzed RNA ligation from the inhibitory effect of excess 2AI. (A)** Increasing concentrations of 2AI inhibits ribozyme-catalyzed ligation in the presence of 10 mM $Mg^{2+}$, which is more than the saturating $Mg^{2+}$ concentration for this ribozyme.[4] **(B)** Increasing concentrations of $Mg^{2+}$ rescue ligation in the presence of 200 mM 2AI. Ligation reactions contained 1.2 µM ribozyme-primer construct, 1.32 µM template, 2.4 µM pre-activated ligator, 200 mM HEPES at pH 8, with the indicated amount of 2AI and $MgCl_2$.

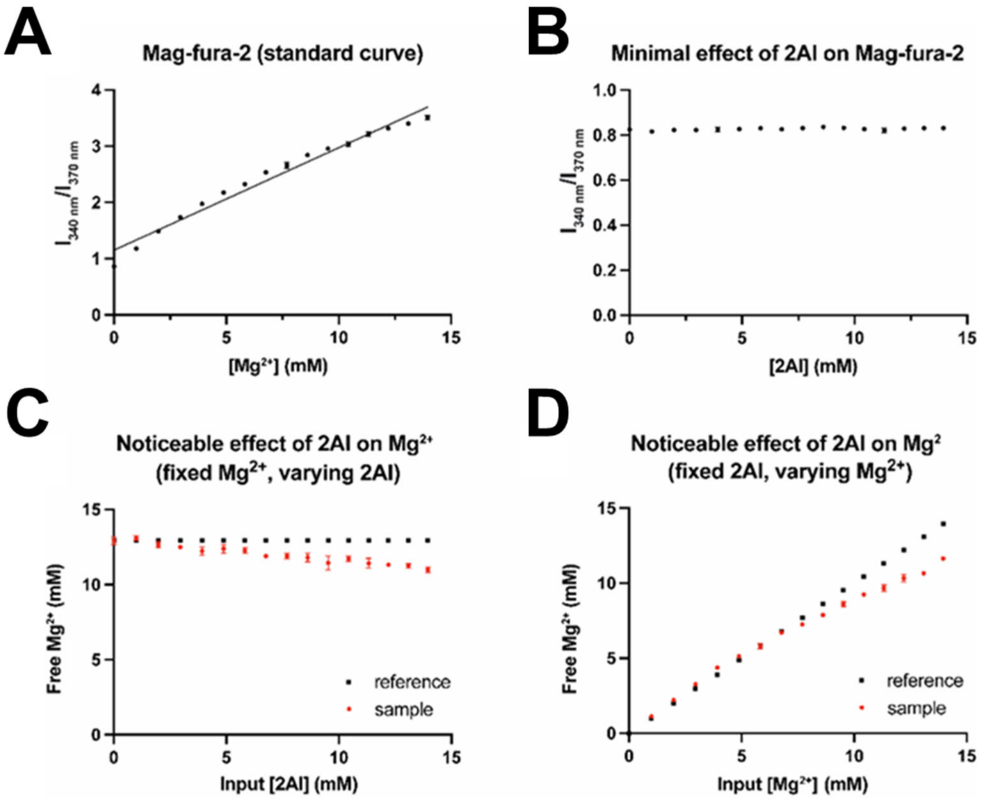


**Fig. S6 A decrease in free $Mg^{2+}$ levels in the presence of 2AI. (A)** Standard curve of Mag-fura-2 fluorescence intensity ($I_{340\ nm}/I_{370\ nm}$) as a function of free $Mg^{2+}$ concentration, with an $R^2$ value of 0.97. **(B)** Mag-fura-2 fluorescence is minimally affected by 2AI up to ~14 mM. At higher 2AI concentrations, deviations from baseline fluorescence are observed, indicating interference with the dye's signal. Therefore, higher concentrations were excluded from the assays shown in (C) and (D). The assay contained 5 µM Mag-fura-2, 200 mM HEPES at pH 8.0, and indicated amounts of 2AI at pH 8.0. **(C)** Free $Mg^{2+}$ concentration decreases with increasing 2AI concentrations (red), as shown by deviations from the reference curve generated in the absence of 2AI (black). This assay was done in the presence of 14 mM $Mg^{2+}$ and varying 2AI concentrations (0-14 mM). **(D)** A deviation from the calibration plot (in black) is observed in the presence of 14 mM 2AI and varying $Mg^{2+}$ concentrations (0–14 mM), indicating reduced free $Mg^{2+}$ levels. Errors are standard deviations of the mean, n = 3 replicates. Assays contained 5 µM Mag-fura-2, 200 mM HEPES at pH 8.0, and the indicated concentrations of 2AI at pH 8.0 and $MgCl_2$.

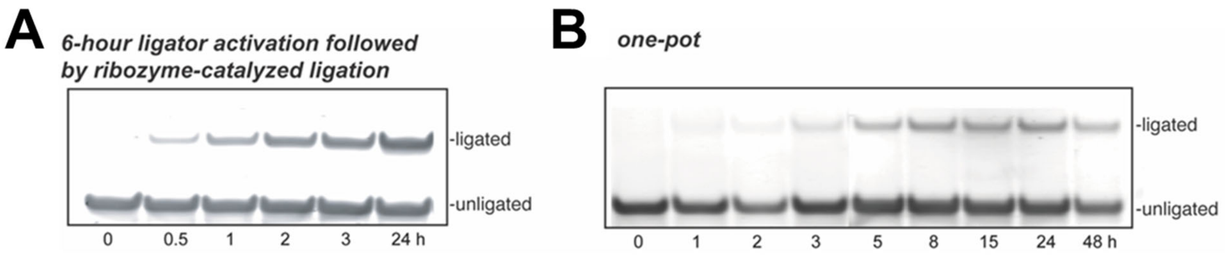


**Fig. S7 Denaturing PAGE depicting the time-course of ribozyme-catalyzed RNA ligation driven by potentially prebiotic oligonucleotide activation.** Ribozyme-catalyzed ligation **(A)** with ligators pre-activated with isocyanide chemistry and **(B)** driven by *in situ* isocyanide activation of ligators were assayed by denaturing PAGE. Reaction conditions: (A) Ribozyme-primer construct and template were added to 2.4 µM isocyanide-activated ligators (250 mM MeNC, 250 mM 2MBA, 250 mM 2AI) in the presence of 200 mM HEPES at pH 8 and 50 mM $MgCl_2$ to a final concentration of 1.2 and 1.32 µM, respectively. (B) 1.2 µM ribozyme-primer construct and 1.32 µM template were directly incubated with 2.4 µM unactivated ligators in the presence of 250 mM MeNC, 250 mM 2MBA, 250 mM 2AI, 200 mM HEPES at pH 8, and 50 mM $MgCl_2$.

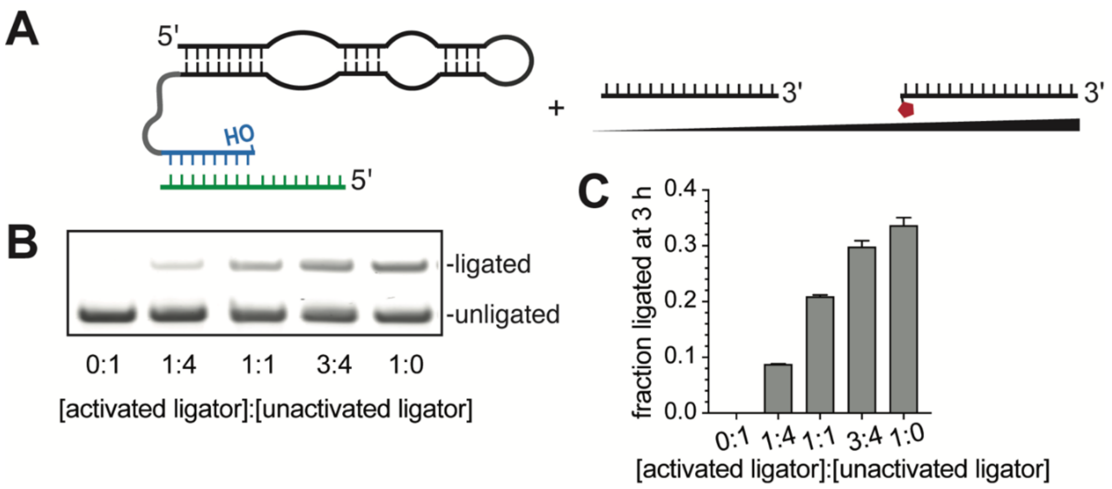


**Fig. S8 Unactivated ligators inhibit ribozyme-catalyzed RNA ligation.** **(A)** Ribozyme-catalyzed RNA ligation in the presence of various ratios of activated to unactivated ligators. **(B)** Ligation reactions containing a mixture of pre-activated and unactivated ligators were analyzed by denaturing PAGE. Ligation was significantly reduced upon increasing the fraction of unactivated ligators. **(C)** Ligation yield gradually decreased from >30% to <10% after 3 h as the ratio of the activated ligator to the unactivated ligator was changed from 1:0 to 1:4. Reactions contained 1.2 µM ribozyme-primer construct, 1.32 µM RNA template, 10 mM $MgCl_2$, 200 mM HEPES at pH 8, and a total of 2.4 µM unactivated and pre-activated 2AI-RNA ligator in varying ratios.

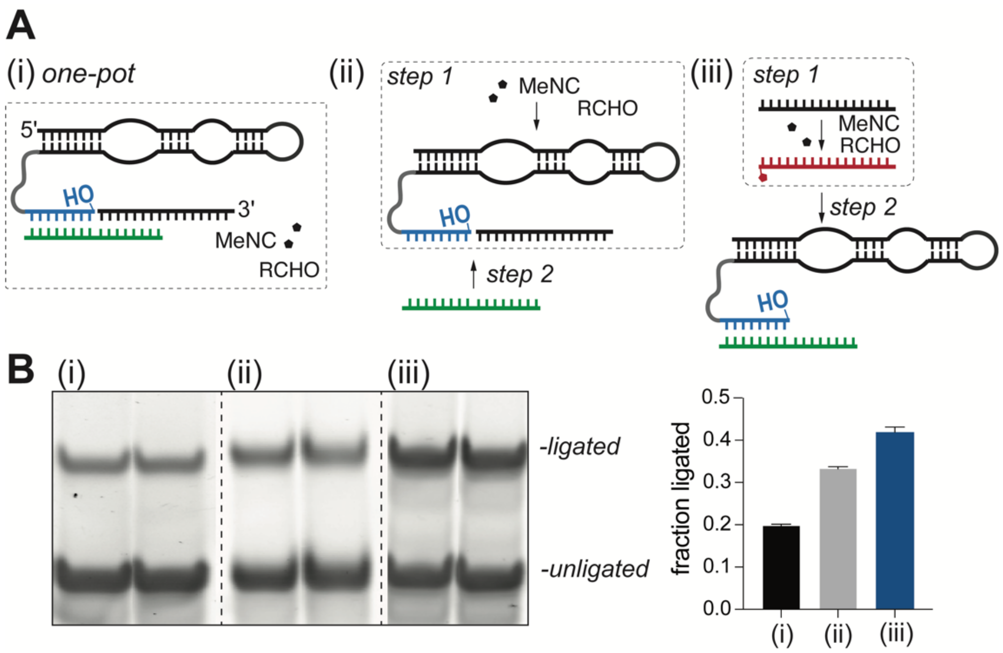


**Fig. S9 The formation of a stable template-ligator duplex hinders ribozyme-catalyzed RNA ligation driven by *in situ* ligator activation. (A)** Ribozyme-catalyzed ligation under three conditions: (i) *in situ* oligomer activation; (ii) *in situ* oligomer activation, where the template is introduced later; and (iii) the addition of ligators pre-activated by isocyanide chemistry. **(B)** Ligation products under the indicated conditions, were assayed and quantified by denaturing PAGE. Error bars represent the standard deviations of the mean, n = 2 replicates. Reaction conditions: (i) 1.2 µM ribozyme-primer construct, 1.32 µM RNA template, 50 mM $MgCl_2$, 200 mM HEPES at pH 8, 2.4 µM unactivated ligator, 250 mM 2MBA, 250 mM MeNC, and 250 mM 2AI. (ii) 1.2 µM ribozyme-primer construct, 50 mM $MgCl_2$, 200 mM HEPES at pH 8, 2.4 µM unactivated ligator, 250 mM 2MBA, 250 mM MeNC, and 250 mM 2AI were allowed to react for 6 h before introducing 1.32 µM RNA template. (iii) 50 mM $MgCl_2$, 200 mM HEPES at pH 8, 2.4 µM unactivated ligator, 250 mM 2MBA, 250 mM MeNC, and 250 mM 2AI were allowed to react for 6 h before introducing 1.32 µM RNA template and 1.2 µM ribozyme-primer construct.

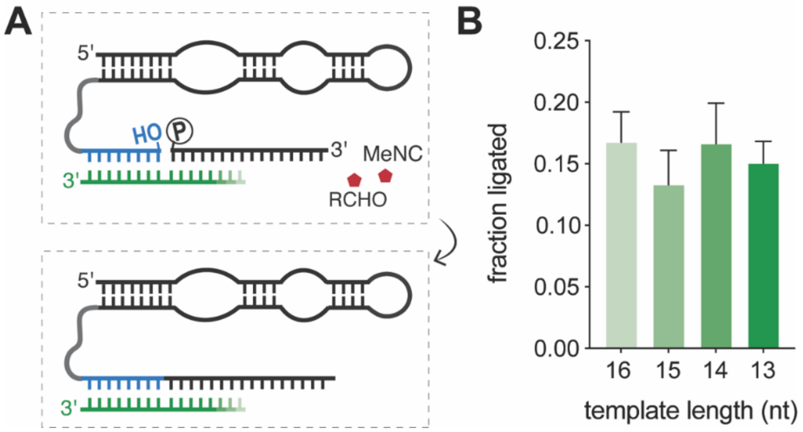


**Fig. S10 Weakening the template-ligator interaction has a modest effect on *in situ* activation-driven, ribozyme-catalyzed RNA ligation. (A)** Shortening the templates by 1-3 nucleotides, which reduces the extent of base-pairing between the ligator and the template, had only a modest effect on **(B)** ligation efficiency. Error bars represent the standard deviations of the mean, n = 2 replicates.

### 3. Supplementary Tables

**Table S1. The yield of 2AI-activated oligonucleotides from corresponding 5′-monophosphorylated oligonucleotides was measured by analytical HPLC following isocyanide chemistry.** 400 mM MeNC, 400 mM 2MBA, and 200 mM 2AI were added to 5 mM oligonucleotides in 200 mM HEPES at pH 8 in the presence of 30 mM $MgCl_2$. The reaction was incubated at room temperature for 6 hours, which was the optimal incubation time as determined previously (Fig. S1B).[1, 5] Prior to HPLC analysis, reagents were removed by Sep-Pak® C18 cartridges. The first two rows of this table use published data from Fig. S5 in reference (5), reused with permission under a Creative Commons Attribution 4.0 International License. Copyright 2023 Ding et al.; Published by Oxford University Press on behalf of *Nucleic Acids Research*.

| Oligonucleotide sequence | [2AI] (mM) | [MeNC] (mM) | [2MBA] (mM) | Activation yield (%) |
|---|---|---|---|---|
| pCA[5] | 200 | 400 | 400 | 90 |
| pCCA[5] | 200 | 400 | 400 | 77 |
| pCGCA[this work] | 200 | 400 | 400 | 87.3 |

**Table S2. Sequences of oligonucleotides used in this study.**

| Name | RNA Sequence (5′ → 3′) |
|---|---|
| Nonenzymatic ligation | |
| 5′-phosphorylated ligator | /5Phos/ ACCACCGCAUUCCGCA |
| Template | GCGGUGGUCAGUCGAG |
| Primer | /5ThioMC6-D/CGCUCGACUG |
| Ribozyme-catalyzed ligation | |
| 5′-phosphorylated ligator | /5Phos/ ACCACCGCAUUCCGCA |
| Template | GCGGUGGUCCUUAGCC |
| Template-1 (15-nt) | CGGUGGUCCUUAGCC |
| Template-2 (14-nt) | GGUGGUCCUUAGCC |
| Template-3 (13-nt) | GUGGUCCUUAGCC |
| Ribozyme-primer construct | GGACAGCGAGCCACUGCGGAAGACCUUAAGAGGUGUAAUUGCUCACCCCGCUG UCCUUUUUUGGCUAAGG |
| Miscellaneous | |
| Unactivated tetramer | /5Phos/ CGCA |
